# The response of zinc transporter gene expression of selected tissues in a pig model of subclinical zinc deficiency

**DOI:** 10.1101/2020.05.14.095919

**Authors:** Daniel Brugger, Martin Hanauer, Johanna Ortner, Wilhelm M. Windisch

## Abstract

This study compared the relative mRNA expression of all mammal zinc (Zn) transporter genes in selected tissues of weaned piglets challenged with short-term subclinical Zn deficiency (SZD). The dietary model involved restrictive feeding (450 g/animal*day^−1^) of a high-phytate diet (9 g/kg) supplemented with varying amounts of zinc from ZnSO_4_*7H_2_O ranging from deficient to sufficient supply levels (total diet Zn: 28.1, 33.6, 38.8, 42.7, 47.5, 58.2, 67.8, 88.0 mg Zn/kg). Total RNA preparations comprised jejunal and colonic mucosa as well as hepatic and nephric tissue. Statistical modelling involved broken-line regression (P ≤ 0.05). ZIP10 and ZIP12 mRNAs were not detected in any tissue and ZnT3 mRNA was only identified in the kidney. All other genes were expressed in all tissues but only a few gene expression patterns allowed a significant (P < 0.0001) fitting of broken-line regression models, indicating homeostatic regulation under the present experimental conditions. Interestingly, these genes could be subcategorized by showing significant turnarounds in their response patterns, either at ~40 or ~60 mg Zn/kg diet (P < 0.0001). In conclusion, the present study showed clear differences in Zn transporter gene expression in response to SZD compared to the present literature on clinical models. We recognized that certain Zn transporter genes were regulated under the present experimental conditions by two distinct homeostatic networks. For the best of our knowledge, this represents the first comprehensive screening of Zn transporter gene expression in a highly translational model to human physiology.

## 1 Introduction

Basic cellular processes are dependent on zinc (Zn) as a structural cofactor of peptides (for example transcription and replication of DNA or maintenance of DNA integrity). In fact, at least 10% of the genes in the human genome encode Zn peptides, highlighting its ubiquitous importance for the mammal organism [1]. In contrast, Zn has a strong toxic potential if the concentration within a biological system exceeds a certain threshold [2]. Therefore, the regulation of Zn uptake, redistribution and excretion within an organism must be tightly controlled.

A complex molecular network that modulates the expression of specific Zn transport peptides to benefit metabolic function under changing dietary and physiological conditions maintains mammalian Zn homeostasis. So far, 24 Zn transporters have been described in mammals mainly based on experiments in rodents and human biopsies. These transporters belong to the solute carrier (SLC) families 30 (ZnT) and 39 (ZIP). Currently, 10 ZnT and 14 ZIP transporter genes have been described [3].

An increasing body of evidence suggests that ZnT and ZIP transporters differ regarding their transport mechanism as well as the direction of Zn transport. The ZnT transporters seem to remove Zn^2+^ from the cytosol, by either facilitating Zn uptake into subcellular compartments or excretion into the extracellular space. In contrast, ZIP transporters increase cytosolic Zn by promoting Zn^2+^ influx from the extracellular space or subcellular compartments, respectively. Zinc transporters also express differences regarding their tissue specificity, subcellular localization as well as the regulative stimuli to which they respond. Furthermore, differences in response patterns of certain transporters have been reported depending on the biological model used for the investigations [3–6]. Liuzzi, Bobo [7] suggested an intestinal-pancreatic axis along which the abundance and activity of certain Zn transporters is regulated on the transcriptional and post-transcriptional level to maintain whole-body Zn homeostasis.

On the one hand, this involves the intestinal regulation of ZIP4 and ZnT1. Current knowledge suggests that ZIP4 is presented on the apical side and ZnT1 on the basolateral side of enterocytes, especially within the jejunum. As a result, these transporters represent the most important active route of luminal Zn transfer into the circulation [7–14]. The regulation of intestinal ZIP4 activity is subject to mechanisms of transcriptional and post-transcriptional regulation. An inadequate dietary Zn supply triggers an increased synthesis of the peptide and its subsequent presentation on the apical plasma membrane. Conversely, body Zn repletion causes ZIP4 endocytosis and breakdown of the protein and its mRNA [15–17]. The amount of ZIP4 mRNA in the enterocyte is a product of changes in its stability and the body Zn status-dependent activity of kruppel-like-factor 4 (KLF4) [18, 19]. In contrast, the expression of *ZnT1* mRNA is a result of the amount of free cytosolic Zn^2+^, which rising concentrations promote the activity of metal-responsive transcription factor 1 (MTF1) [13, 20, 21]. Therefore, *ZnT1* gene expression is no marker for the body Zn status but the cytosolic Zn^2+^ levels in certain tissues.

Most endogenous Zn losses occur in the gastrointestinal tract [22], mainly via exogenous pancreatic secretions of Zn-metallopeptides (e.g. zymogens like carboxypeptidase A and B) [23]. The differential uptake of Zn in pancreatic acinar cells and the excretion back into the circulation are due to the activity of ZIP5 and ZnT1 at the basolateral membrane. A Zn deficiency reduces the ZIP5-dependent uptake from the circulation and at the same time promotes the activity of ZnT1, presumably as a measure to bring Zn back into the periphery for the benefit of other tissues. Zinc repletion and excess promote the opposite response [7, 16, 24]. The transporter that loads Zn into zymogen granules of pancreatic acinar cells is ZnT2. A systemic Zn deficiency decreases the pancreatic ZnT2 abundance, which has associated with a decrease in the Zn concentrations in these granules. Otherwise, a systemic Zn overload triggers the opposite reaction, as a measure in favor of an increased excretion of excess Zn quantities from the system into the gastrointestinal tract [25].

These aforementioned molecular changes in the intestine and exogenous pancreas nicely reflect the organisms attempts to adjust gastrointestinal absorption and endogenous Zn losses in times of deficiency and oversupply, respectively, and directly correspond to classical data on quantitative Zn fluxes in ^65^Zn labelled rats fed varying dietary Zn concentrations [26].

Subclinical zinc deficiency is probably the most common form of zinc malnutrition in humans and animals [27]. However, it has not yet been thoroughly investigated. In fact, many of the original *in vivo* studies on the role of zinc in metabolism and nutrition included control groups expressing symptoms of clinical zinc deficiency. Under such conditions, however, the compensation capacities of the metabolism, and in particular the mobilizable Zn pools in the skeleton and the soft tissues, are exhausted [28–31]. This results in the quite unspecific set of visible symptoms of Zn malnutrition (e.g. growth depression, anorexia, developmental disorders, tissue necrosis etc. in pigs [32]) and, as a result, an increased background noise from metabolic measurements. In addition, this represents the endpoint in adapting metabolic processes to an ongoing dietary zinc deficiency, which can reduce the informative value of quantitative measurements of Zn-homeostatic mechanisms at the level of the whole organism.

To promote translational research on SZD, we have developed an experimental model to promote this phenotype in pigs [33]. It allows high-resolution analyses of the kinetics and dynamics of zinc in the growing pig organism, from the level of the quantitative metabolism to the subcellular level. At the same time, it does not promote any change in the health status of the animals based on continuous veterinary surveillance. The previous findings revealed the adjustment of the quantitative zinc metabolism to a short-term (8d), finely graded reduction in the alimentary zinc intake. Based on this, we were able to derive the gross zinc requirement under the given experimental conditions at 60 mg / kg diet [33]. Already in this early phase of zinc deficiency, various pathophysiological reactions at the metabolic and subclinical level became evident. These included a reduction in pancreatic digestive capacity [34] and cardiac redox capacity [35]. The latter was accompanied by an increased need for the detoxification of reactive oxygen species and the activation of pro-apoptotic signaling pathways when the alimentary zinc supply was ~40 mg / kg diet and below for a period of 8 days [35]. Furthermore, we were able to show that certain tissues that are important for the acute survival of the developing organism (heart, skeletal muscle, thymus, mesenteric lymph nodes, pancreas) partly or fully replenished their initially depleted zinc concentrations at the expense of other organs or even accumulated Zn above the level of the control group. The critical threshold for these compensatory reactions was also ~40 mg Zn / kg diet [36]. Overall, these findings suggested two independent Zn-homeostatic regulatory pathways that act in this early phase of developing zinc deficiency: 1) the regulation of the absorption, redistribution and excretion of zinc for the maintenance of whole-body Zn homeostasis (critical threshold: ~60 mg / kg diet for 8d) and 2) the compensation of increased oxidative stress and perhaps inflammation in certain tissues as a result of an ongoing nutritional zinc deficiency (critical threshold; ~40 mg / kg diet for 8d).

So far, no study has dealt with the qualitative and quantitative expression of all known Zn transporter genes in a large animal model. In addition, current data sets on Zn transporters are mostly limited to very specific roles related to biomedical research questions, and only a handful of studies looked at the situation under non-clinical conditions in and between tissues in response to finely graded differences in dietary Zn supply. Finally, the regulation of Zn transporters under conditions of SZD is yet unclear. We believe that the comparative analysis of the Zn transporter gene expression in and between tissues allows a deeper insight into the adaptation of the body to different Zn supply levels. Such a dataset could also be suitable to test our hypothesis of two different homeostatic regulation pathways in the early stages of a developing Zn deficiency. The present study therefore examined the expression of the known ZnT and ZIP genes in potentially important tissues of zinc homeostasis (jejunal and colonic mucosa, liver, kidney). To the best of our knowledge, this dataset represents the first comprehensive examination of ZnT and ZIP gene expression in the translational model of the pig. Furthermore, it appears to be the first investigation of zinc transporter genes in the context of such a high-resolution dose-response study *in vivo*.

## 2 Material and Methods

This animal study was evaluated and approved by the animal welfare officer of the faculty TUM School of Life Sciences Weihenstephan, Technical University of Munich, and further approved and registered by the responsible animal welfare authorities (District Government of Upper Bavaria, Federal State of Bavaria: case number 55.2.1.54-2532.3.63-11).

The study design as well as communication of material, methods and results comply to the ARRIVE Guidelines [37]. The dietary model applied for this investigation was carefully developed to allow in-depth physiological research in a large translational animal model with a minimum of animals (n = 6 replicates / treatment group). For further details on the dietary model and the statistical consideration during study preparation see Brugger, Buffler [33] and the subsection on statistical analyses, respectively.

### 2.1 Animals and diets

This study applied the experimental SZD model originally proposed by Brugger, Buffler [33], in which weaned piglets are adapted to a phytate-rich basal diet and subsequently treated with varying dietary Zn supplementation for 8d. Therefore, a total of forty-eight fully weaned piglets from six litters (hybrids of German Large White x Land Race x Piétrain, 8 animals per litter, 50% male-castrated, 50% female, initial average body weight 8.5 ±0.27 kg, 28 d of age, supplier: pig farm of Christian Hilgers (Germany) were individually housed. To ensure full body Zn stores at day one of the experimental period, a basal diet (based on corn and soybean meal, dietary phytate (InsP6) concentration 9 g/kg) with a native Zn concentration of 28.1 mg/kg was supplemented with 60 mg Zn/kg from analytical-grade ZnSO_4_ * 7H_2_O to adjust the total dietary Zn at a sufficient level of 88.0 mg/kg diet. This diet was provided to all animals ad libitum during an acclimatization phase of 14d prior to the onset of the experiment. Subsequently, all animals were assigned to eight dietary treatment groups in a complete randomized block design. Blocking parameters comprised live weight, litter mates and sex (50% male-castrated, 50% female), thereby yielding a good standardization of body development and genetic background between treatment groups. The treatment groups were fed restrictively the same basal diet as during the acclimatization period (450 g/d representing the average *ad libitum* feed intake at the last day of acclimatization) but with varying dietary Zn concentrations spanning the range from deficient to sufficient dietary supply levels in high resolution. This was achieved by differential supplementation of analytical-grade ZnSO_4_ * 7H_2_O (+0, +5, +10, +15, +20, +30, +40, +60 mg added Zn/kg diet; analyzed dietary Zn contents: 28.1 ± 0.24, 33.6 ± 0.56, 38.8 ± 0.82, 42.7 ± 0.48, 47.5 ± 0.97, 58.2 ± 0.29, 67.8 ± 1.87, 88.0 ± 1.65 mg Zn/kg diet). The group receiving 88.0 mg Zn/kg diet served as control, because it represented the feeding situation during the acclimatization phase from which the dietary Zn contents for all other groups were gradually reduced. The total experimental period during which the varying zinc feeding occurred lasted a total of 8d. Figure 1 shows a scheme of the study time-course.

**Figure 1.**
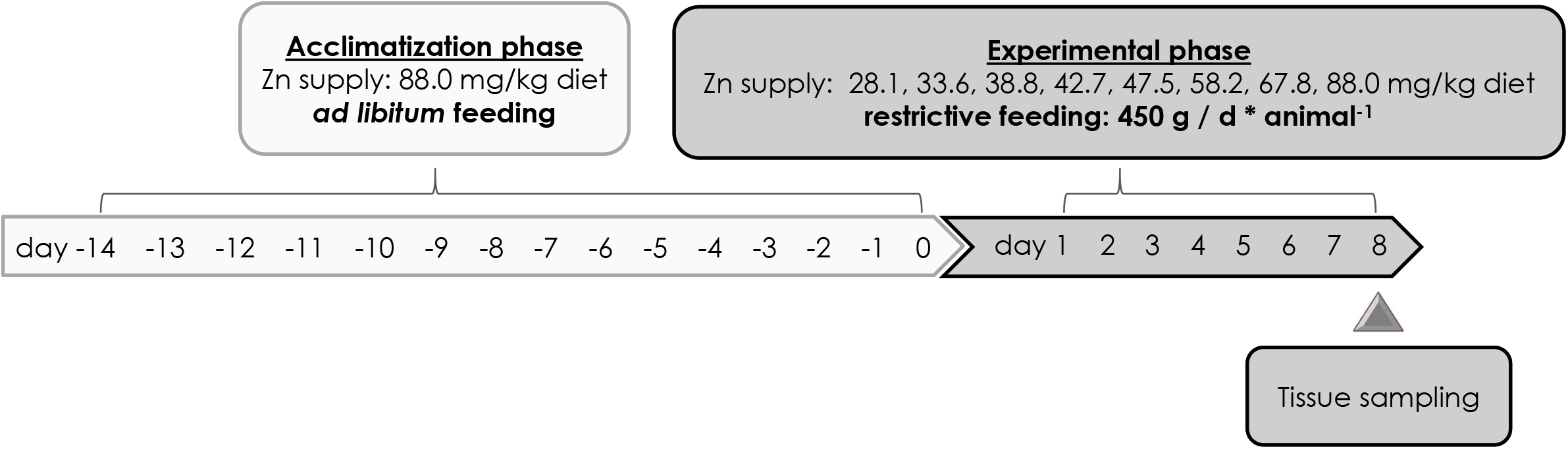
Time-course of the study. Piglets were assigned to eight treatment groups in a complete randomized block design. Blocking parameters comprised live weight, litter mates and sex (50% male-castrated, 50% female). During a 14d acclimatization period, all animals were fed a diet with sufficient dietary Zn supply (+60 mg Zn/kg diet resulting in 88 mg/kg diet) *ad libitum*. During a subsequent experimental phase of 8d, treatment groups were fed restrictively (450 g/d) the same basal diet as during the acclimatization period but with varying dietary Zn concentrations (+0, +5, +10, +15, +20, +30, +40, +60 mg Zn/kg diet resulting in 28.1, 33.6, 38.8, 42.7, 47.5, 58.2, 67.8, 88.0 mg Zn/kg diet). Analytical grade *ZnSO_4_*7H_2_O* was used for varying Zn supplementation. In the end of experimental d8, all animals were killed by exsanguination under anesthesia (azaperone and ketamine) without fasting to obtain tissue samples. All diets met or exceeded published recommendations for the feeding of weaned piglets according to NRC [21] except for Zn. d, day(s); g, gram, kg, kilogram; mg, milligram, ZnSO_4_*7H_2_O, zinc sulfate heptahydrate; Zn, zinc.

The basal diet was designed to meet all the recommendations of the National Research Council regarding the feeding of weaned piglets except for Zn [38]. Table 1 presents detailed information on the composition and ingredients of the basal diet. Analytical Zn recovery of added Zn from the experimental diets as an indicator of the mixing precision was literally 100% as shown by a highly significant slope of 0.99 mg increase in total analyzed dietary Zn per mg Zn addition to the diets (P < 0.0001, R^2^ = 1.00, data not shown). All diets were pelletized at 70°C with steam to stabilize feed particle size distribution, improve feed hygiene and deactivate native phytase activity originating from plant raw components.

**Table 1.**
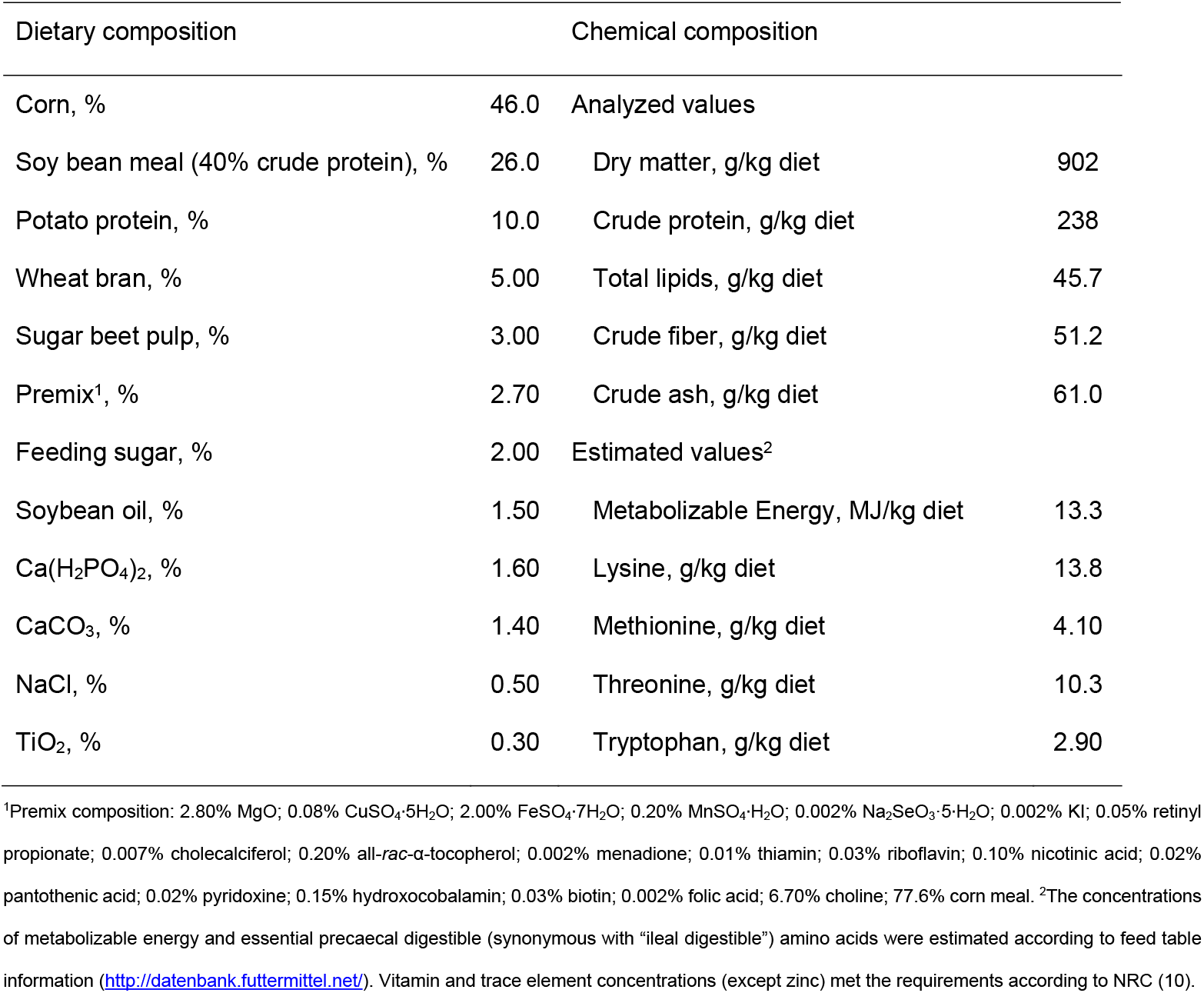
Composition as well as concentrations of metabolizable energy and crude nutrients of the basal diet [33]

All animals had access to drinking water (tap water) *ad libitum* during all times of this study and were subject to continuous veterinary surveillance. The zinc background in drinking water was analyzed regularly to ensure negligible background levels.

### 2.2 Sampling conditions

Diet samples were collected and processed as described previously [33]. At experimental day 8, all animals were killed by exsanguination under anesthesia (azaperone and ketamine) without fasting and tissue samples were taken from jejunal and colonic mucosa as well as liver and kidney. Tissue samples for gene expression analyses were immediately incubated in RNAlater (Life Technologies GmbH) overnight and subsequently stored at −80°C according to the manufacturer’s instructions.

### 2.3 Analyses of dry matter and total zinc concentration in diets

Analyses of dry matter and Zn in diets occurred as described earlier [33]. Each experimental diet was sampled in triplicate and milled through a 0.5 mm screen. Subsequently, each of the triplicates was weighed in triplicate for chemical extraction procedures, producing n = 9 data points for nutrient and Zn concentrations of each individual experimental diet). Zn concentrations were measured by atomic absorption spectrometry (AAS) (NovAA 350; Analytik Jena AG) applying a certified AAS Zn standard material (Merck 109953, Merck Millipore) after microwave wet digestion (Ethos 1, MLS GmbH) with 65% HNO_3_, 30% H_2_O_2_ and Type I ultrapure water.

### 2.4 Gene expression analysis

Primer design, assay quality control and chemical procedures (total RNA extraction, reverse transcription, quantitative PCR (qPCR)) were performed as described earlier[33, 35]. Purity (measured with the NanoDrop 2000 system, Thermo Scientific) and integrity (measured with the Experion system, Biorad) of all total RNA extracts from all tissues met or exceeded the minimum thresholds necessary for gene expression profiling applying qPCR methodology[39]. Primer pairs (Eurofins Scientific) were designed with Primer Blast [40] for the potential reference transcripts glyceraldehyde 3-phosphate dehydrogenase (GAPDH), β-glucuronidase (GUSB), histone H3 (H3), ubiquitin C (UBC), β-actin (ACTB) and divalent metal transporter 1 (DMT1) as well as the target transcripts SLC30 (ZnT) 1, 2, 3, 4, 5, 6, 7, 8, 9, 10 and SLC39 (ZIP) 1, 2, 3, 4, 5, 6, 7, 8, 9, 10, 11, 12, 13, 14, based on published porcine sequence information [41] (Supplementary Tables 1 and 2). All oligonucleotides bind to homologous regions of respective transcripts to amplify the pool of potential transcript variants within one reaction. Reference gens were selected for each individual tissue by applying the RefFinder tool for the respective complete Ct datasets from each tissue [42]. This tool combines the major computational approaches for reference evaluation (geNorm, Normfinder, BestKeeper, comparative Delta-Ct method) to compare and rank the tested genes. In this way we identified the following suitable reference genes: *ZnT5, ZnT6* and *ZIP9* in the jejunum, *DMT1, ZIP1* and *ZnT5* in the colon, *DMT1, ZIP7* and *ZIP13* in the liver as well as *ZnT4, ZnT5* and *ZnT6* in the kidney. The 2^−ΔΔCt^ method [43] was used to normalize the gene expression data, because the determination of the amplification efficiency revealed comparable values between 95% and 100% of applied RT-qPCR assays. A detailed description of the procedure for the determination of amplification efficiency has been provided earlier [33].

*ZIP10* and *ZIP12* transcripts were not detected in any of the porcine tissues examined within the present study. These assays amplify sequences, which appear to be highly conserved between mammal species. Therefore, murine brain and liver cDNA preparations were used for testing.

### 2.5 Statistical analyses

Data analysis was performed with SAS 9.4 (SAS Institute Inc.) applying the procedure NLMIXED to estimate linear broken-line regression models (*y = a + bx + cx*) based on independent group means relative to dietary Zn concentration (*n = 8*). The decision for linear broken-line models over non-linear models was made by following the approach proposed by McDonald [44] by which the goodness-of-fit of linear vs. polynomial models is statistically compared using F-statistics. In case of our dataset we found that applying non-linear models over the linear broken-line models yielded no significant increase in the quality of the curve-fitting. Furthermore, using single data points from individual animals instead of the group mean values was not advisable in light of the present experimental design because the imbalance in the ratio of X (eight dietary treatment groups) to Y (six response values per treatment group) coordinates would have caused a severe overestimation of the degrees of freedom and, therefore, false results. Broken-line regression is an iterative procedure to estimate a potential statistical threshold (breakpoint) within non-linear data sets above and below which a significant difference in the response behavior of a certain parameter to the dietary treatment is evident [45]. If no significant breakpoint in parameter response could be estimated from a certain data set, a linear regression model was tested instead (y= a + bx) (procedure REG). It is noteworthy, that those gene expression patterns, which yielded no significant broken-line regression model, also did not fit significant linear models. Only significant regression models were applied for data presentation and interpretation in the present manuscript. A threshold of P ≤ 0.05 was considered to indicate statistical significance. All *2*^−ΔΔCt^ gene expression values [43] were presented as x-fold differences compared to a relative mRNA abundance of 1.0 (not regulated) within the control group (88.0 mg Zn/kg diet).

The justification of the total sample size for the present experiment was done prior to the study by power analysis with the software package G*Power 3.1.9.6 [46] assuming a two-factor model (8 treatment groups, 6 experimental blocks) including interactions (treatment*block) and a strong biological effect. Based on this analysis it has been concluded that forty-eight animals (n = 6 replicates/treatment group) in a completely randomized block design are sufficient to meet the generally accepted minimum statistical power of 1 – β = 0.8 [47]. The correctness of these assumptions were confirmed by estimating the power of the applied regression models and associate T-statistics on regression parameters, which met in any case the necessary minimum of 1-β = 0.8.

## 3 Results

All animals remained in good health throughout the whole trial. There were no signs of clinical Zn deficiency in swine (for example growth retardation, anorexia as earlier reported by Tucker and Salmon [32]) evident at any time [33]. The experimental model introduced finely-graded adaptions in Zn status parameters as well as the Zn concentrations in the tissues under study (Supplementary Figure 1). This data has been presented earlier [36] and will therefore not been described in more detail in the present manuscript.

### 3.1 Tissue specificity of *ZnT* and *ZIP* transcripts in weaned piglets challenged with finely graded differences in zinc supply status

Table 2 highlights the qualitative expression pattern of analyzed transcripts within respective tissues. Most of the analysed *ZnT* and *ZIP* transcripts were abundant within the tissues examined in the present study. This excludes *ZIP10* and *ZIP12*, which were not expressed in any of the tissues as well as *ZnT3*, which was only recognised within the kidney. Testing *ZIP10* and *ZIP12* assays in murine liver and brain cDNA preparations yielded positive results and excluded technical problems to be the cause of negative results derived in porcine cDNA from jejunum, colon, liver and kidney, respectively. Some transcripts (*ZnT5, ZnT6* and *ZIP9* in jejunum, *ZIP1* and *ZnT5* in colon, *ZIP7* and *ZIP13* in liver as well as *ZnT4, ZnT5* and *ZnT6* in kidney) were expressed in such a highly stable manner over treatment groups that they served as reference genes for data normalisation (based on data analyses using earlier published statistical approaches [48]).

**Table 2:** Qualitative expression pattern of ZnT and ZIP genes within the jejunum, colon, liver and kidney of weaned piglets fed diets with different Zn concentrations for 8d^1^.

| Transcript | Jejunum | Colon | Liver | Kidney |
| --- | --- | --- | --- | --- |
| ZnT1 | ✓ | ✓ | ✓ | ✓ |
| ZnT2 | ✓ | ✓ | ✓ | ✓ |
| ZnT3 | N/A | N/A | N/A | ✓ |
| ZnT4 | ✓ | ✓ | ✓ | ✓<sup>R</sup> |
| ZnT5 | ✓<sup>R</sup> | ✓<sup>R</sup> | ✓ | ✓<sup>R</sup> |
| ZnT6 | ✓<sup>R</sup> | ✓ | ✓ | ✓<sup>R</sup> |
| ZnT7 | ✓ | ✓ | ✓ | ✓ |
| ZnT8 | ✓ | ✓ | ✓ | ✓ |
| ZnT9 | ✓ | ✓ | ✓ | ✓ |
| ZnT10 | ✓ | ✓ | ✓ | ✓ |
| ZIP1 | ✓ | ✓<sup>R</sup> | ✓ | ✓ |
| ZIP2 | ✓ | ✓ | ✓ | ✓ |
| ZIP3 | ✓ | ✓ | ✓ | ✓ |
| ZIP4 | ✓ | ✓ | ✓ | ✓ |
| ZIP5 | ✓ | ✓ | ✓ | ✓ |
| ZIP6 | ✓ | ✓ | ✓ | ✓ |
| ZIP7 | ✓ | ✓ | ✓<sup>R</sup> | ✓ |
| ZIP8 | ✓ | ✓ | ✓ | ✓ |
| ZIP9 | ✓<sup>R</sup> | ✓ | ✓ | ✓ |
| ZIP10 | N/A | N/A | N/A | N/A |
| ZIP11 | ✓ | ✓ | ✓ | ✓ |
| ZIP12 | N/A | N/A | N/A | N/A |
| ZIP13 | ✓ | ✓ | ✓<sup>R</sup> | ✓ |
| ZIP14 | ✓ | ✓ | ✓ | ✓ |

Notes: <sup>1</sup>The applied dietary Zn concentrations were 28.1, 33.6, 38.8, 42.7, 47.5, 58.2, 67.8 and 88.0 mg Zn/kg. <sup>R</sup>evaluated as a suitable reference gene using published mathematical procedures [48]. N/A, transcript not detected in respective tissue sample; SLC, solute carrier; ZIP1 to 14, SLC family 39 members 1 to 14; ZnT1 to 10, SLC family 30 members 1 to 10.

### 3.2 Effects of varying dietary zinc supply on the relative *ZnT* and *ZIP* transcript abundance in examined porcine tissues of weaned piglets challenged with finely graded differences in zinc supply status

Many transcripts recognized within the jejunum, colon, liver and kidney of growing piglets showed significant dietary thresholds in response to changes in dietary Zn supply. This was evident by significant breakpoint parameter estimates (*P* ≤ 0.05 for X and Y intercepts of respective breakpoints). The only exceptions were *ZIP2* and *ZIP3* in colonic tissue as well as the candidate genes that served as reference genes for data normalisation within respective tissues. Significant dietary thresholds either lay at ~40 or ~60 mg Zn/kg diet, respectively. However, the slopes of the respective segments within many broken-line regression models were not significant and coefficients of determination of respective models were low (R^2^). Subsequently, only models expressing at least one significant slope over changes in dietary Zn supply are described within figures and tables.

Figure 2 presents the broken-line response of jejunal Zn transporter gene expression as affected by varying dietary Zn supply. Table 3 presents the corresponding statistical measures of the respective regression curves. Above significant dietary thresholds of 57.1, 62.3, 38.8, 41.6, 62.6 and 52.3 mg Zn/kg diet (*P* < *0.0001*, respectively) jejunal *ZIP5* and *ZIP11* significantly increased or decreased, respectively, in response to changes in dietary Zn (*P ≤ 0.05*, respectively) whereas *ZIP1* and *ZIP13* did not change in a significant manner. Below these thresholds, the relative mRNA abundance of *ZIP1* and *ZIP13* significantly increased whereas *ZIP11* significantly decreased with further reduction in dietary Zn supply (*P ≤ 0.001, ≤ 0.05* and ≤ *0.0001*, respectively). The *ZIP5* gene expression did not change significantly with stepwise decrease in dietary Zn concentration below its respective breakpoint.

**Figure 2.**
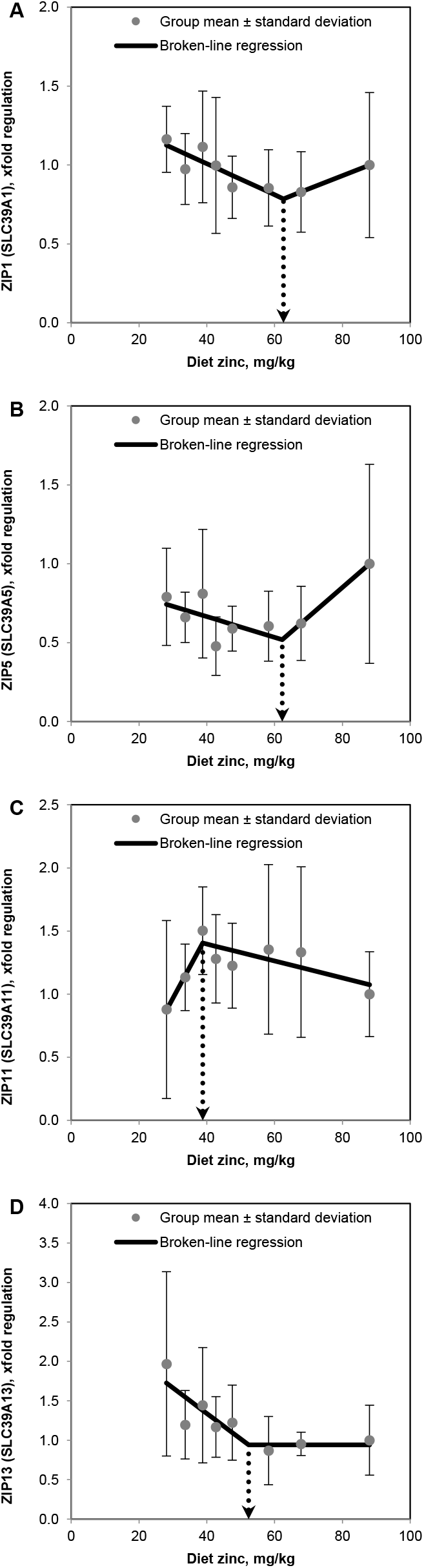
Response of relative jejunal gene expression of ZIP1 (A), ZIP5 (B), ZIP11 (C) and ZIP13 (D) in weaned piglets fed diets with different zinc concentrations for 8 d (see Table 3 for detailed information on the statistical measures of the respective regression models). Values are arithmetic means ± SDs, *n = 6*. d, day; diet zinc, dietary zinc; xfold, difference in the gene expression value according to Livak and Schmittgen [27] compared to a relative mRNA abundance of 1.0 in the control group (88.0 mg Zn/kg diet); ZIP1, 5, 11, 13, solute carrier (SLC) family 39 members 1, 5, 11, 13.

**Table 3.** Broken-line regression analysis of relative jejunal gene expression (xfold) of ZIP1, 5, 11 and 13 in weaned piglets fed diets with different zinc concentrations for 8d^1^.

| | Regression model | Breakpoint | Slopes | $R^2$ |
| --- | --- | --- | --- | --- |
| ZIP1 | $y = 1.40 + b_1x \text{ for } x \leq X_B$ | $X_B \ 62.6^{***} \pm 5.07$ | $b_1 \ -0.01^{**} \pm 0.002$ | 0.75 |
| | $y = 0.26 + b_2x \text{ for } x \geq X_B$ | $Y_B \ 0.79^{***} \pm 0.05$ | $b_2 \ 0.008 \pm 0.004$ | |
| ZIP5 | $y = 0.93 + b_1x \text{ for } x \leq X_B$ | $X_B \ 62.3^{***} \pm 5.38$ | $b_1 \ -0.007 \pm 0.003$ | 0.70 |
| | $y = -0.64 + b_2x \text{ for } x \geq X_B$ | $Y_B \ 0.52^{***} \pm 0.07$ | $b_2 \ 0.02^* \pm 0.006$ | |
| ZIP11 | $y = -0.52 + b_1x \text{ for } x \leq X_B$ | $X_B \ 38.8^{***} \pm 0.02$ | $b_1 \ 0.05^{***} \pm 0.009$ | 0.79 |
| | $y = 1.66 + b_2x \text{ for } x \geq X_B$ | $Y_B \ 1.40^{***} \pm 0.05$ | $b_2 \ -0.007^* \pm 0.002$ | |
| ZIP13 | $y = 2.63 + b_1x \text{ for } x \leq X_B$ | $X_B \ 52.3^{***} \pm 6.05$ | $b_1 \ -0.03^* \pm 0.01$ | 0.75 |
| | $y = 0.94 \text{ for } x \geq X_B$ | $Y_B \ 0.94^{***} \pm 0.09$ | N/A | |

Notes: <sup>1</sup>The applied dietary zinc concentrations were 28.1, 33.6, 38.8, 42.7, 47.5, 58.2, 67.8, and 88.0 mg Zn/kg. Broken-line regression models were estimated based on independent arithmetic group means relative to dietary zinc concentration ( $n = 8$ ). Parameter estimates are presented as means ± SEs to indicate the precision of estimation. $P \leq 0.05$ was considered to be significant. \*, \*\*, \*\*\* indicate $P \leq 0.05$ , 0.001, 0.0001, respectively. $b_1$ , slope of the broken-line regression curves over dietary zinc doses $\leq X_B$ ; $b_2$ , slope of the broken-line regression curves over dietary zinc doses $> X_B$ ; SLC, solute carrier; xfold, difference in the gene expression value according to Livak and Schmittgen [43] compared to a relative mRNA abundance of 1.0 in the control group (88.0 mg Zn/kg diet); $X_B$ , X intercept of the breakpoint in the parameter response; $Y_B$ , Y intercept of the breakpoint in the parameter response; ZIP1, 5, 11, 13, SLC family 39 members 1, 5, 11, 13.

Figure 3 presents the broken-line response of colonic Zn transporter gene expression as affected by varying dietary Zn supply. Table 4 presents the corresponding statistical measures of the respective regression curves. Relative mRNA abundance of *ZnT4, ZnT9, ZIP4, ZIP5, ZIP7, ZIP11* and *ZIP13* showed significant breakpoints in response to a finely graded reduction in dietary Zn concentration at 60.6, 63.9, 59.6, 39.0, 42.7, 44.8 and 68.3 mg Zn/kg diet, respectively (*P ≤ 0.0001*, respectively). Above the respective dietary thresholds, *ZIP4, ZIP5* and *ZIP7* plateaued in response to changes in dietary Zn. These genes significantly increased their relative expression levels in response to further reduction in dietary Zn below these breakpoints (*P ≤ 0.001, ≤ 0.001* and ≤ *0.05* for *ZIP4, ZIP5* and *ZIP7*, respectively). On the contrary, colonic *ZnT4* and *ZIP11* significantly increased (*P ≤ 0.05* and ≤ *0.001*, respectively) whereas *ZnT9* and *ZIP13* significantly decreased (*P ≤ 0.05*, respectively) with reduction in dietary Zn concentration from 88.0 mg Zn/kg to their respective breakpoints. Below these dietary thresholds, *ZnT4* and *ZIP11* significantly decreased (*P ≤ 0.001* and ≤ *0.05*, respectively) whereas *ZnT9* and *ZIP13* did not change significantly.

**Figure 3.**
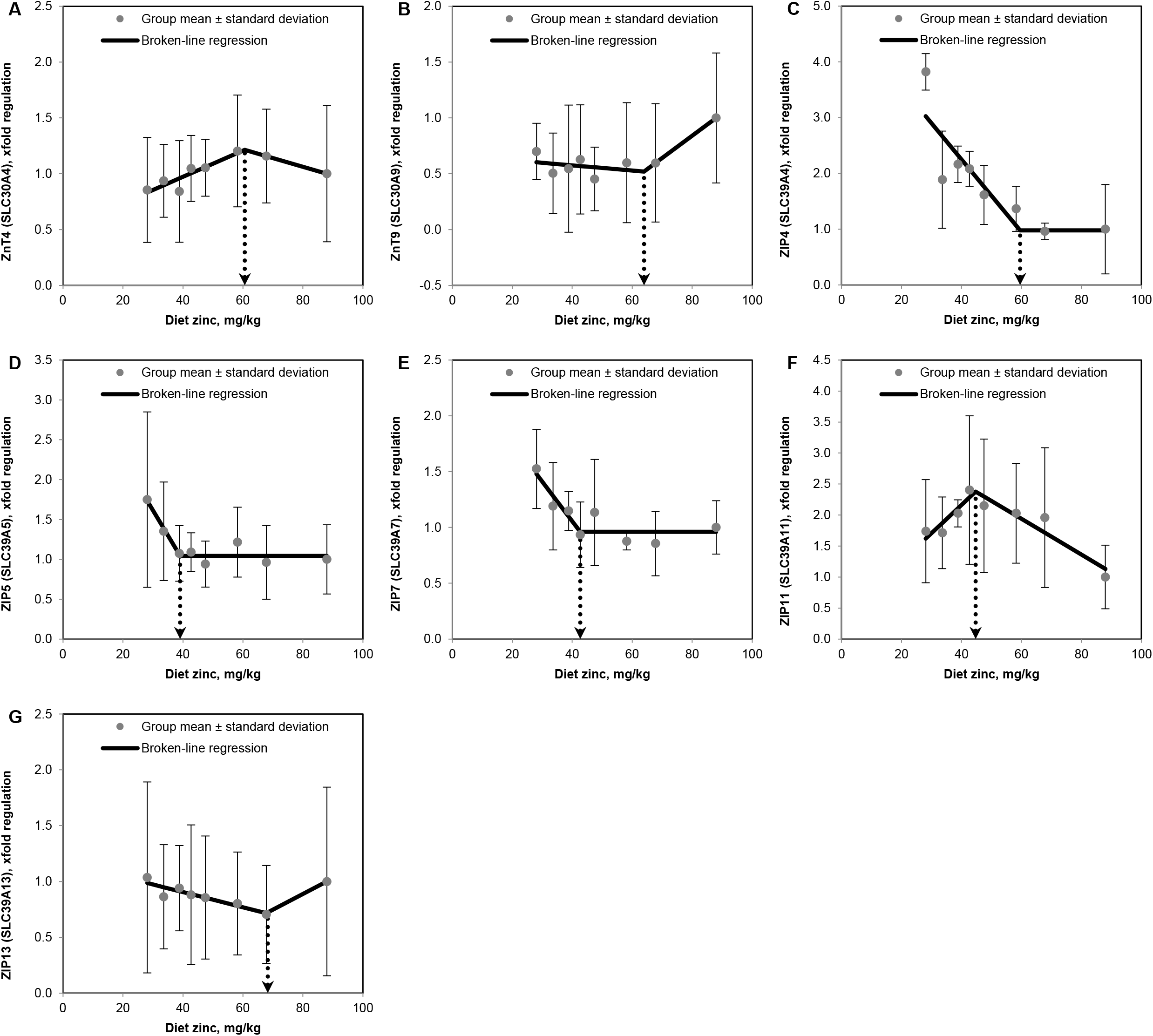
Response of relative colonic gene expression of ZnT4 (A), ZnT9 (B), ZIP4 (C), ZIP5 (D), ZIP7 (E), ZIP11 (F) and ZIP13 (G) in weaned piglets fed diets with different zinc concentrations for 8 d (see Table 4 for detailed information on the statistical measures of the respective regression models). Values are arithmetic means ± SDs, n = 6. d, day; diet zinc, dietary zinc; xfold, difference in the gene expression value according to Livak and Schmittgen [27] compared to a relative mRNA abundance of 1.0 in the control group (88.0 mg Zn/kg diet); ZnT4, 9, solute carrier (SLC) family 30 members 4, 9; ZIP4, 5, 7, 11, 13, SLC family 39 members 4, 5, 7, 11, 13.

**Table 4.** Broken-line regression analysis of relative colonic gene expression (xfold) of ZnT4 and 9 as well as ZIP4, 5, 7, 11 and 13 in weaned piglets fed diets with different zinc concentrations for 8 d^1^.

| | Regression model | Breakpoint | Slopes | $R^2$ |
| --- | --- | --- | --- | --- |
| ZnT4 | $y = 0.50 + b_1x \text{ for } x \leq X_B$ | $X_B \ 60.6^{***} \pm 4.01$ | $b_1 \ 0.01^{**} \pm 0.002$ | 0.85 |
| | $y = 1.69 + b_2x \text{ for } x \geq X_B$ | $Y_B \ 1.21^{***} \pm 0.04$ | $b_2 \ -0.008^* \pm 0.003$ | |
| ZnT9 | $y = 0.67 + b_1x \text{ for } x \leq X_B$ | $X_B \ 63.9^{***} \pm 4.82$ | $b_1 \ -0.002 \pm 0.003$ | 0.82 |
| | $y = -0.75 + b_2x \text{ for } x \geq X_B$ | $Y_B \ 0.52^{***} \pm 0.06$ | $b_2 \ 0.02^* \pm 0.005$ | |
| ZIP4 | $y = 4.85 + b_1x \text{ for } x \leq X_B$ | $X_B \ 59.6^{***} \pm 7.14$ | $b_1 \ -0.06^{**} \pm 0.02$ | 0.76 |
| | $y = 0.98 \text{ for } x \geq X_B$ | $Y_B \ 0.98^* \pm 0.29$ | N/A | |
| ZIP5 | $y = 3.51 + b_1x \text{ for } x \leq X_B$ | $X_B \ 39.0^{***} \pm 1.32$ | $b_1 \ -0.06^{**} \pm 0.01$ | 0.90 |
| | $y = 1.04 \text{ for } x \geq X_B$ | $Y_B \ 1.04^{***} \pm 0.04$ | N/A | |
| ZIP7 | $y = 2.48 + b_1x \text{ for } x \leq X_B$ | $X_B \ 42.7^{***} \pm 2.50$ | $b_1 \ -0.04^* \pm 0.008$ | 0.81 |
| | $y = 0.96 \text{ for } x \geq X_B$ | $Y_B \ 0.96^{***} \pm 0.05$ | N/A | |
| ZIP11 | $y = 0.34 + b_1x \text{ for } x \leq X_B$ | $X_B \ 44.8^{***} \pm 2.43$ | $b_1 \ 0.04^* \pm 0.01$ | 0.87 |
| | $y = 3.67 + b_2x \text{ for } x \geq X_B$ | $Y_B \ 2.38^{***} \pm 0.09$ | $b_2 \ -0.03^{**} \pm 0.005$ | |
| ZIP13 | $y = 1.18 + b_1x \text{ for } x \leq X_B$ | $X_B \ 68.3^{***} \pm 5.3$ | $b_1 \ -0.007^{***} \pm 0.001$ | 0.86 |
| | $y = -0.26 + b_2x \text{ for } x \geq X_B$ | $Y_B \ 0.71^{***} \pm 0.04$ | $b_2 \ -0.01^* \pm 0.004$ | |

Notes: <sup>1</sup>The applied dietary zinc concentrations were 28.1, 33.6, 38.8, 42.7, 47.5, 58.2, 67.8, and 88.0 mg Zn/kg. Broken-line regression models were estimated based on independent arithmetic group means relative to dietary zinc concentration ( $n = 8$ ). Parameter estimates are presented as means ± SEs to indicate the precision of estimation. $P \leq 0.05$ was considered to be significant. \*, \*\*, \*\*\* indicate $P \leq 0.05$ , 0.001, 0.0001, respectively. $b_1$ , slope of the broken-line regression curves over dietary zinc doses $\leq X_B$ ; $b_2$ , slope of the broken-line regression curves over dietary zinc doses $> X_B$ ; SLC, solute carrier; xfold, difference in the gene expression value according to Livak and Schmittgen [43] compared to a relative mRNA abundance of 1.0 in the control group (88.0 mg Zn/kg diet); $X_B$ , X intercept of the breakpoint in the parameter response; $Y_B$ , Y intercept of the breakpoint in the parameter response; ZIP4, 5, 7, 11, 13, SLC family 39 members 4, 5, 7, 11, 13 ZnT4, 9, solute carrier (SLC) family 30 members 4, 9.

Figure 4 presents the broken-line response of hepatic Zn transporter gene expression as affected by varying dietary Zn supply. Table 5 presents the corresponding statistical measures of the respective regression curves. Gene expression patterns of *ZnT4, ZnT6, ZnT8, ZIP1* and *ZIP14* exhibited significant changes in their response to varying dietary Zn supply at breakpoints of 48.4, 38.8, 57.3, 47.5 and 42.7 mg Zn/kg diet, respectively (*P ≤ 0.0001*). Above the respective dietary thresholds, *ZnT4, ZnT8, ZIP1* and *ZIP14* did not change significantly in response to changes in dietary Zn supply whereas *ZnT6* significantly decreased directly to a reduction in dietary Zn from 88.0 mg Zn/kg diet to the respective breakpoint (*P ≤ 0.001*). On the contrary, *ZnT4* and *ZnT6* significantly decreased (*P ≤ 0.001* and ≤ *0.05*, respectively) whereas *ZnT8, ZIP1* and *ZIP14* significantly increased (*P ≤ 0.05, ≤ 0.05* and ≤ *0.0001*, respectively) with reduction in dietary Zn concentration below the respective dietary thresholds.

**Figure 4.**
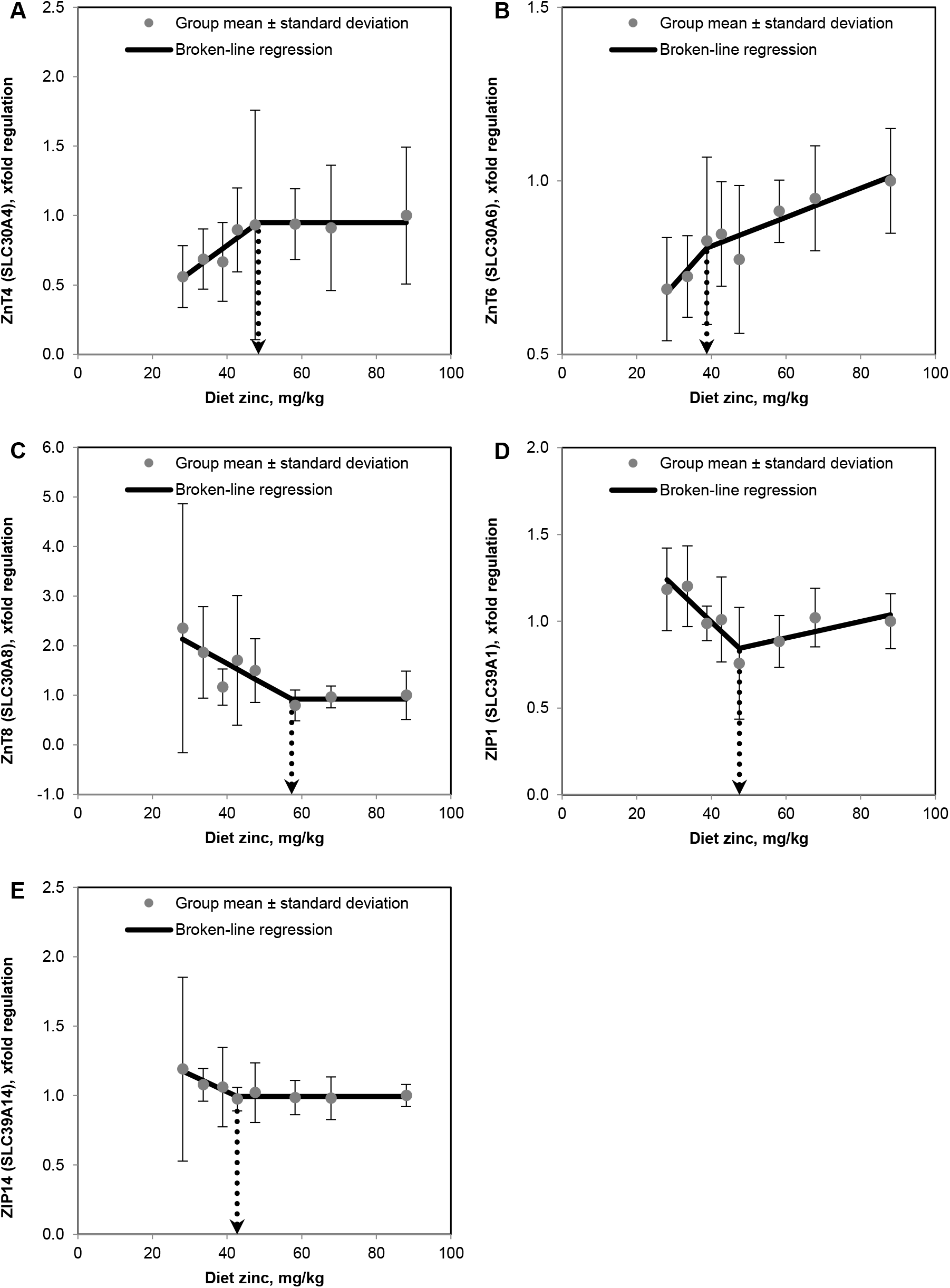
Response of relative hepatic gene expression of ZnT4 (A), ZnT6 (B), ZnT8 (C), ZIP1 (D) and ZIP14 (E) in weaned piglets fed diets with different zinc concentrations for 8 d (see Table 5 for detailed information on the statistical measures of the respective regression models). Values are arithmetic means ± SDs, n = 6. d, day; diet zinc, dietary zinc; xfold, difference in the gene expression value according to Livak and Schmittgen [27] compared to a relative mRNA abundance of 1.0 in the control group (88.0 mg Zn/kg diet); ZnT4, 6, 8 solute carrier (SLC) family 30 members 4, 6, 8; ZIP1, 14, SLC family 39 members 1, 14.

**Table 5.**
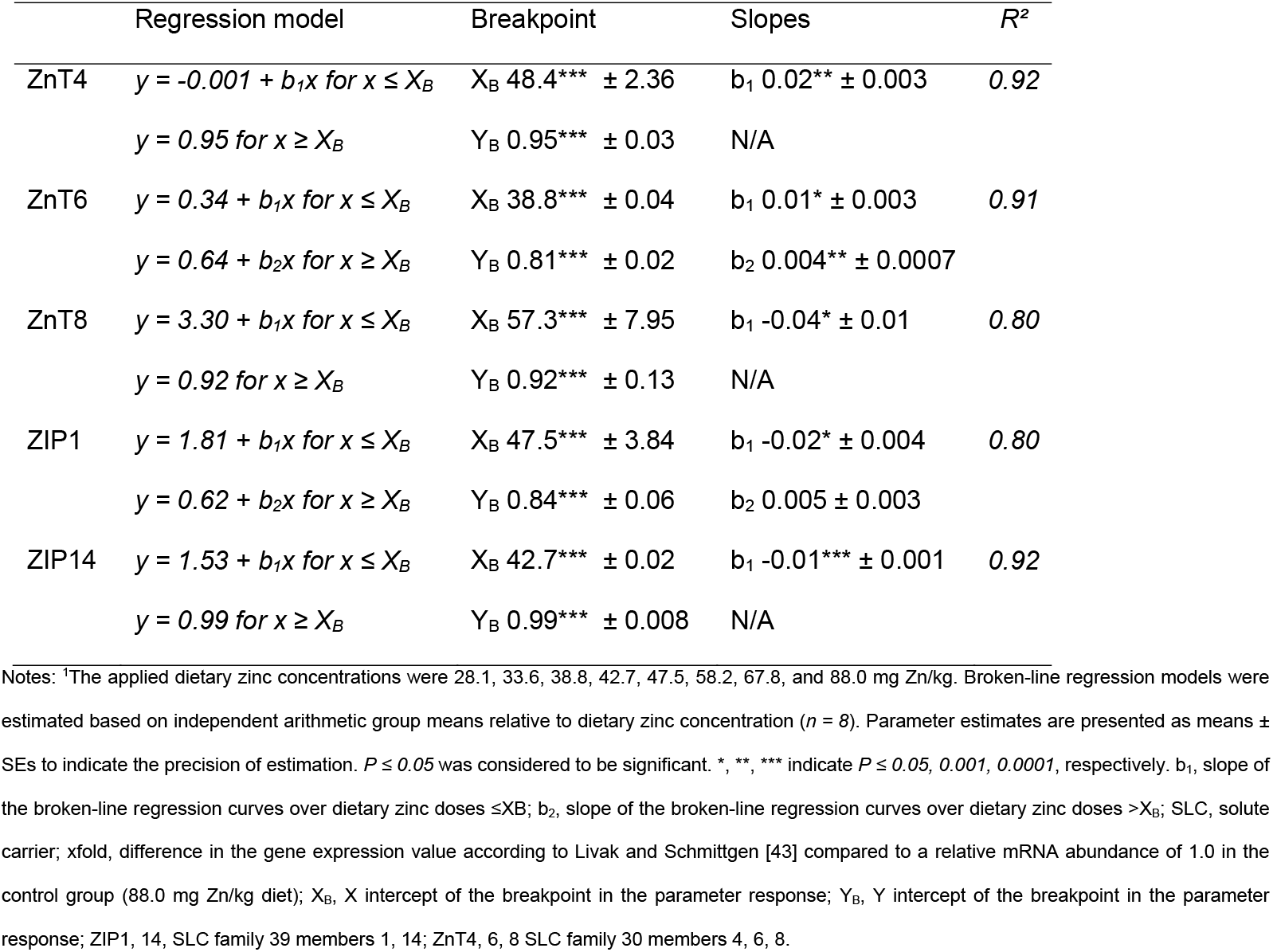
Broken-line regression analysis of relative hepatic gene expression (xfold) of ZnT4, 6 and 8 as well as ZIP1 and 14 in weaned piglets fed diets with different zinc concentrations for 8 d^1^.

Figure 5 presents the broken-line response of nephric Zn transporter gene expression as affected by varying dietary Zn supply. Table 6 presents the corresponding statistical measures of the respective regression curves. Gene expression of *ZnT1, ZnT3, ZnT7* and *ZIP4* changed significantly around dietary thresholds of 70.4, 42.6, 35.2 and 41.9 mg Zn/kg diet (*P* < *0.0001*, respectively). All these genes plateaued in response to a reduction of dietary Zn concentration from 88.0 mg/kg to the respective breakpoints. Further reduction in dietary Zn below these thresholds promoted a significant increase of *ZnT3, ZnT7* and *ZIP4 (P ≤ 0.05, ≤ 0.0001* and ≤ *0.05*, respectively) as well as a significant decrease of *ZnT1* gene expression (*P ≤ 0.05*).

**Figure 5.**
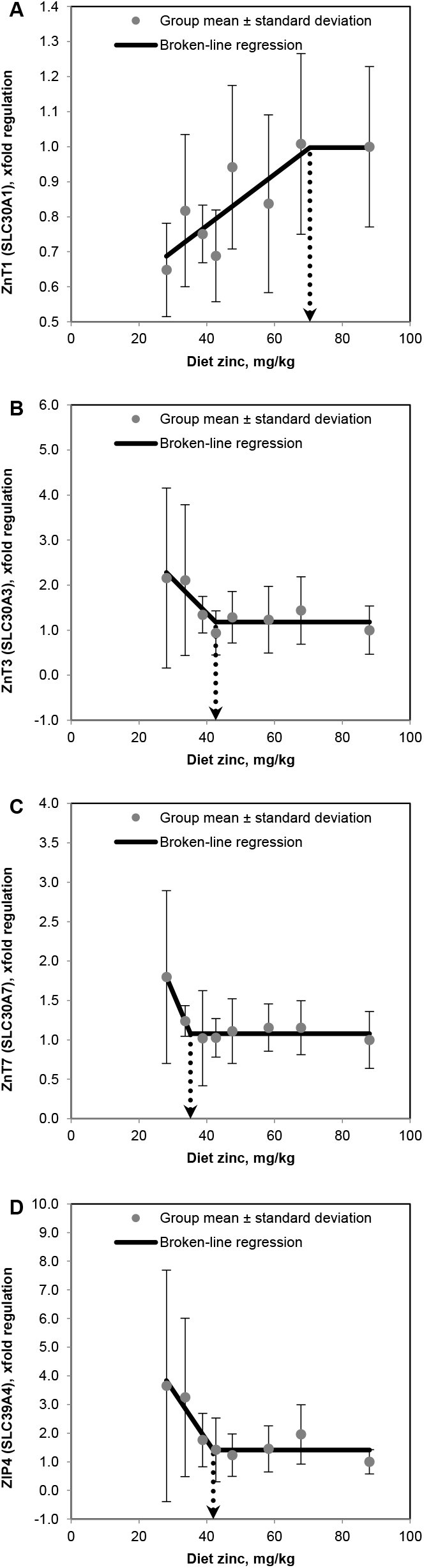
Response of relative nephric gene expression of ZnT1 (A), ZnT3 (B), ZnT7 (C) and ZIP4 (D) in weaned piglets fed diets with different zinc concentrations for 8 d (see Table 6 for detailed information on the statistical measures of the respective regression models). Values are arithmetic means ± SDs, n = 6. d, day; diet zinc, dietary zinc; xfold, difference in the gene expression value according to Livak and Schmittgen [27] compared to a relative mRNA abundance of 1.0 in the control group (88.0 mg Zn/kg diet); ZnT1, 3, 7, solute carrier (SLC) family 30 members 1, 3, 7, ZIP4, SLC family 39 member 4.

**Table 6.** Broken-line regression analysis of relative nephric gene expression (xfold) of ZnT1, 3 and 7 as well as ZIP4 in weaned piglets fed diets with different zinc concentrations for 8 d^1^.

| | Regression model | Breakpoint | Slopes | $R^2$ |
| --- | --- | --- | --- | --- |
| ZnT1 | $y = 0.48 + b_1x \text{ for } x \leq X_B$ | $X_B \ 70.4^{***} \pm 9.84$ | $b_1 \ 0.007^* \pm 0.002$ | 0.75 |
| | $y = 1.00 \text{ for } x \geq X_B$ | $Y_B \ 1.00^{***} \pm 0.05$ | N/A | |
| ZnT3 | $y = 4.40 + b_1x \text{ for } x \leq X_B$ | $X_B \ 42.6^{***} \pm 3.40$ | $b_1 \ -0.08^* \pm 0.02$ | 0.82 |
| | $y = 1.18 \text{ for } x \geq X_B$ | $Y_B \ 1.18^{***} \pm 0.08$ | N/A | |
| ZnT7 | $y = 4.65 + b_1x \text{ for } x \leq X_B$ | $X_B \ 35.2^{***} \pm 0.76$ | $b_1 \ -0.10^{***} \pm 0.01$ | 0.95 |
| | $y = 1.08 \text{ for } x \geq X_B$ | $Y_B \ 1.08^{***} \pm 0.02$ | N/A | |
| ZIP4 | $y = 8.78 + b_1x \text{ for } x \leq X_B$ | $X_B \ 41.9^{***} \pm 2.26$ | $b_1 \ -0.18^* \pm 0.04$ | 0.89 |
| | $y = 1.41 \text{ for } x \geq X_B$ | $Y_B \ 1.41^{***} \pm 0.13$ | N/A | |

Notes: <sup>1</sup>The applied dietary zinc concentrations were 28.1, 33.6, 38.8, 42.7, 47.5, 58.2, 67.8, and 88.0 mg Zn/kg. Broken-line regression models were estimated based on independent arithmetic group means relative to dietary zinc concentration (n = 8). Parameter estimates are presented as means ± SEs to indicate the precision of estimation. $P \leq 0.05$ was considered to be significant. \*, \*\*\* indicate $P \leq 0.05$ , 0.0001, respectively. b<sub>1</sub>, slope of the broken-line regression curves over dietary zinc doses $\leq X_B$ ; b<sub>2</sub>, slope of the broken-line regression curves over dietary zinc doses $> X_B$ ; SLC, solute carrier; xfold, difference in the gene expression value according to Livak and Schmittgen [43] compared to a relative mRNA abundance of 1.0 in the control group (88.0 mg Zn/kg diet); X<sub>B</sub>, X intercept of the breakpoint in the parameter response; Y<sub>B</sub>, Y intercept of the breakpoint in the parameter response; ZIP4, SLC family 39 member 4; ZnT1, 3, 7, SLC family 30 members 1, 3, 7.

Figure 6 summarizes the aforementioned breakpoint estimates from Tables 3, 4, 5 and 6 including respective standard errors of the statistical estimation. Across tissues, observed breakpoints of gene expression response to varying dietary Zn supply scattered around mean values of 42.9 ±2.08 or 63.1 ±6.19 mg Zn/kg diet, respectively.

**Figure 6.**
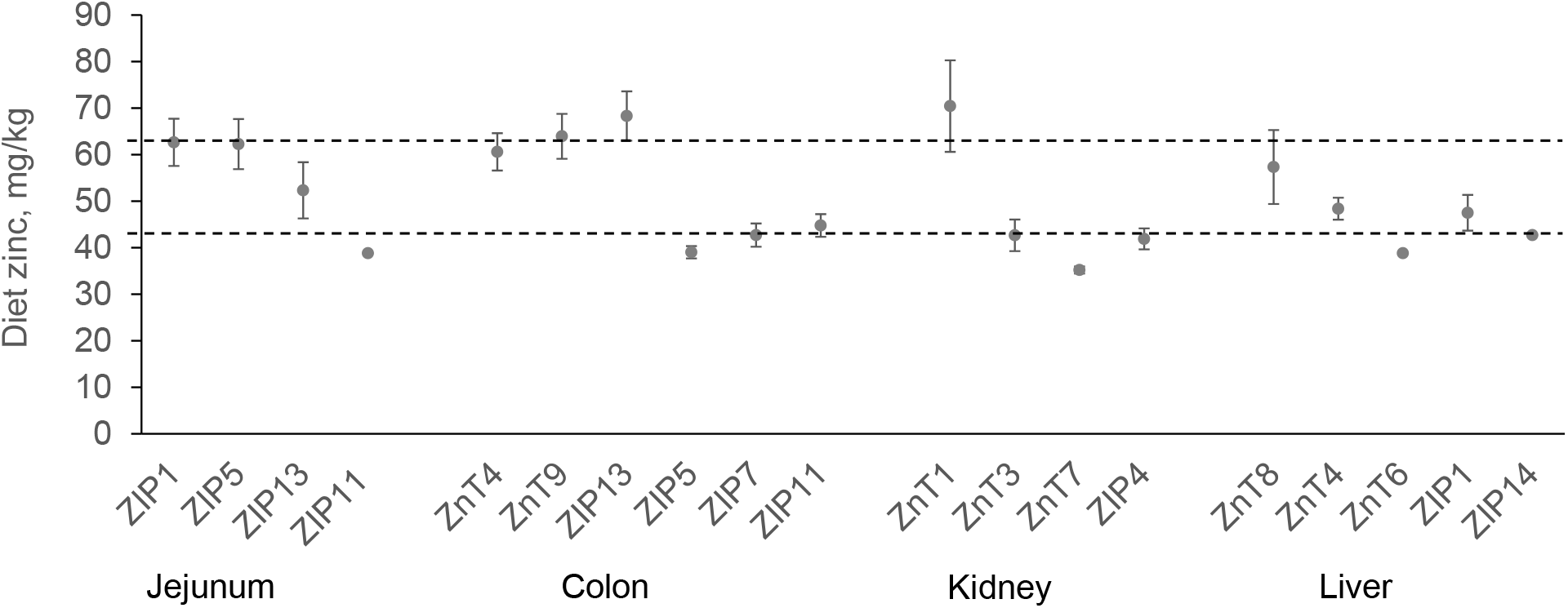
Comparative presentation of dietary breakpoint estimates for ZIP and ZnT gene expression patterns in jejunum, colon, liver and kidney of weaned piglets fed diets with different zinc concentrations for 8 d (see Figures 2, 3, 4 and 5 as well as Tables 3, 4, 5 and 6 for the respective regression models). Across tissues, breakpoints scattered around dietary thresholds of either 42.9 ±2.08 or 63.1 ±6.19 mg Zn/kg diet, respectively, marked by horizontal dashed lines. Error bars indicate standard errors of breakpoint estimation. Each breakpoint estimate was highly significant (P < 0.0001 in any case). The presentation of breakpoints for each tissue was ordered in a way to present the higher before the lower values. Breakpoints were obtained by broken-line regression analysis over each eight treatment means, which were calculated based on single values of each six animals per group fed restrictively (450 g/d) a diet with varying dietary Zn concentrations (28.1, 33.6, 38.8, 42.7, 47.5, 58.2, 67.8, 88.0 mg Zn/kg diet) in a complete randomized block design. Analytical grade ZnSO_4_*7H_2_O was used for varying Zn supplementation. ZnT, solute carrier (SLC) family 30, ZIP, SLC family 39, Zn, zinc.

## 4 Discussion

We investigated the expression of ZnT (SLC30) and ZIP (SLC39) transporter genes in weaned piglets exposed to different Zn concentrations in the diet (28.1 to 88.0 mg Zn/kg). The tissues examined were jejunal and colonic mucosa, liver and kidney, as the play key roles in in the uptake, (re)distribution and excretion of Zn in the body [23]. The gene expression of the pancreas was not investigated for technical reasons. The experimental setup included both sexes and the results of the present study were comparable between them.

### 4.1 Specificity of ZnT and ZIP family member gene expression for selected porcine tissues

Most transcripts were present in all examined tissues, which seems to agree with the available literature (summarized in [3–6]). The only exceptions were *ZIP10* and *ZIP12*, which were not recognized at all. In addition, *ZnT3* mRNA was only expressed in nephric tissue.

RNA sequencing experiments in humans and *Drosophila* confirmed *ZIP12* gene expression only in the brain [49, 50]. In addition, studies on its functional genomics and proteomics have highlighted its role in neuronal development and function [51, 52] and in the schizophrenic brain [53]. Hence, the absence of *ZIP12* cDNA in any tissue from the present study agrees well with these findings. Otherwise, *ZIP10* transcription has been identified in various tissues, including those investigated in the present study [49, 50]. Aside from its role in cancer progression, its function as a zinc transporter within the renal collecting duct system has been studied in various species [54–56], highlighting its possible role in reabsorption of Zn^2+^ from primary urine. Hence, the question arises why we could not identify its mRNA signature in any of our samples, including those from the kidney? Our *ZIP10* qPCR assay amplifies an mRNA sequence that is conserved in mammals, including rodents and humans. It worked successfully with murine brain and liver preparations (data not shown). Furthermore, we used a representative cross-section of each porcine kidney for the total RNA preparations. We therefore conclude that the lack of *ZIP10* transcripts in our tissue samples was not due to technical bias and that the growing pig does not express this gene in respective tissues under the given experimental conditions. However, future studies should apply microdissection of different parts of the porcine kidney to confirm our results.

*ZnT3* mRNA was found in brain, adipose tissue, pancreatic beta-cells, epithelial cells, testis, prostate and retina [57]. To date, most studies have focused on its role in transferring cytosolic Zn into synaptic vesicles [58]. Interestingly, this was also the case in gastrointestinal enteric neurons from pigs [59, 60]. The lack of *ZnT3* gene expression in pig jejunum and colon from the present study may be due to the fact that these samples represented the mucosa. To our knowledge, no study has yet investigated *ZnT3* gene expression in the livers and kidneys of pigs. Therefore, this could be the first report on the lack of *ZnT3* gene expression in the liver and its simultaneous detection in the kidney of the growing pig, at least under SZD conditions.

### 4.2 Response of *ZnT* and *ZIP* family member gene expression to dietary zinc

It has previously been shown that some Zn transporter genes respond to deficient dietary Zn supply (*ZnT1, ZnT2, ZnT4, ZnT5, ZnT6, ZIP4, ZIP10*) [13, 18, 55, 61]. We have confirmed this for *ZnT1* (kidney), *ZnT4* (colon, liver), *ZnT6* (liver) and *ZIP4* (colon, kidney) but not for *ZnT2, ZnT5* and *ZIP10*. In addition, other transcripts also responded to deficient dietary Zn supply, which apparently have not yet been reported, including *ZnT3* (kidney), *ZnT7* (kidney), *ZnT8* (liver), *ZnT9* (colon), *ZIP1* (jejunum, liver), *ZIP5* (jejunum, colon), *ZIP7* (colon), *ZIP11* (jejunum, colon), *ZIP13* (jejunum, colon) and *ZIP14* (liver). Most earlier studies used clinical Zn deficiency models. Therefore, our data seem to highlight differences in the adaption of Zn transporter gene expression to short-term SZD. Clinical Zn deficiency is associated with many secondary metabolic events during which tissue integrity can be compromised [32]. This reduces the resolution of measurements due to increased background noise. Finally, clinical Zn deficiency is the endpoint in physiological adaption to body Zn depletion. Hence, early response patterns have already changed, which could explain why some results from the present short-term study were not described earlier. Several of the *ZnT* and *ZIP* gene expression patterns showed significant breakpoints in response to diet Zn. However, only the reaction of the genes shown in the figures and tables allowed a significant curve fitting. We assume that only these genes were directly regulated by changes in the dietary Zn supply. On the contrary, the Zn transporter genes with only significant breakpoints may have just adapted indirectly to the changes in Zn fluxes triggered by these key transporters and are therefore not discussed further in this manuscript.

Genes showing a significant dose-response to dietary Zn could be divided into two groups, with statistical breakpoints at either ~40 or ~60 mg Zn/kg diet. The gross Zn requirement under given conditions was earlier estimated at ~60 mg Zn/kg diet [33]. We therefore assume that *ZnT* and *ZIP* genes, with a statistical breakpoint of ~60 mg Zn/kg diet, are involved in maintaining whole-body Zn homeostasis. A prominent example was colonic *ZIP4* gene expression, which plateaued in groups receiving ≥60 mg Zn/kg diet. A gradual decrease in dietary Zn below this threshold induced a linear upregulation of *ZIP4* gene expression. In fact, this response pattern correlates significantly with earlier data on the apparently digested feed Zn under the present experimental conditions (*r = −0,91; P = 0.002*; data not shown) [33]. This reflects the classic concept of Zn homeostasis, according to which the organism increases its Zn absorption capacity at the gut barrier during periods of Zn malnutrition [26]. It also agrees with earlier published work in which ZIP4 was identified as the main active Zn transporter at the apical membrane of enterocytes [16, 18, 24]. Our data confirms the role of this gene within the homeostatic network that controls Zn levels in the body and particularly Zn absorption in pigs and other mammals.

Regarding genes with a statistical breakpoint at ~40 mg Zn/kg diet, we postulate that they may be involved in regulation of tissue Zn in connection with redox and immune functions. We have shown earlier under the present experimental conditions that the heart muscle of Zn-deficient piglets restored its initially depleted total Zn concentration. This was probably due to an up-regulation of the transporters that import Zn from the circulation into the tissue. It occurred as a reaction to increased cardiac oxidative stress during SZD in groups fed ≤40 mg Zn/kg diet [35]. Therefore, breakpoints of Zn transporter genes at ~40 mg Zn/kg diet could indicate similar events in other tissues. A prominent example is the hepatic ZIP14. It was previously identified as an acute-phase protein that transports circulating Zn and non-transferrin iron into the liver during times of systemic inflammation [62, 63]. In the present study, its gene expression increased in piglets fed ≤42.7 mg Zn/kg diet. Given the previous cardiac data, this could suggest that these animals developed an inflammatory state during SZD. Earlier published data support this hypothesis by showing an inverse correlation between systemic inflammatory activity and Zn status in the elderly [64]. The functional background has recently been reviewed [65]. Indeed, we currently lack data on the adaption of stress and inflammatory pathways in the tissues studied. These must be collected in future studies to allow a functional correlation to the regulation and activity of certain Zn transporters.

It must be considered that the threshold of ~40 mg Zn/kg diet does not represent a minimum dietary Zn concentration above which there are no adverse effects on the redox metabolism. Rather, it seems to be related to the maximum tolerable bone Zn depletion during our experiment. In fact, animals that received ≤43 mg Zn/kg diet showed a reduction in bone Zn between ~20-25% under given experimental conditions [33]. This represents an exhaustion of the mobilizable skeletal Zn fraction, which was previously shown in ^65^Zn-labelled rats [66–68]. A continuation of the study >8 d until mobilizable body Zn stores were finally depleted, would most likely have increased this threshold over time until it had reached the previously defined gross Zn requirement [33].

### 4.3 ZIP4 and ZnT1 expression in the intestine and kidney under SZD conditions

ZIP4 and ZnT1 are the active route of Zn transfer from the intestinal lumen to the circulation [7–14]. Previous studies suggest that *ZIP4* transcription directly responds to the status of whole-body Zn homeostasis through KLF4 activity [18, 19]. Colonic *ZIP4* expression has already been discussed. Its breakpoint reflects the gross Zn requirement under given test conditions (60 mg/kg) [33] and confirms regulation for the benefit of whole-body Zn homeostasis. Interestingly, we did not find any significant response of jejunal *ZIP4* gene expression. The jejunum is generally regarded as the main site of Zn absorption [69]. We therefore expected a significant upregulation of *ZIP4* transcription in the jejunal before the colonic mucosa, like has been shown in mice fed low Zn diets [7, 9, 16, 70]. This was not the case, which again highlights clear differences between subclinical and clinical Zn deficiency. Given the response of colonic *ZIP4* transcription to changes in dietary Zn supply and its positive correlation with apparent Zn absorption, we assume that the large intestine is the main site of Zn acquisition under SZD conditions. This has already been suggested by other authors who found peaks in the colonic expression of Zn-responsive genes under conditions of mild Zn deficiency in adult rats [21, 71]. In addition, there have been reports of a significant contribution of caecal and colonic Zn absorption in times of impaired Zn acquisition from the small intestine [72, 73]. We have previously shown a decrease in pancreatic digestive capacity under the present experimental conditions [34]. We therefore conclude that the main site of Zn absorption may have shifted to the large intestine in favor of Zn-dependent digestive enzymes to stabilize the already impaired protein digestion. This needs to be checked in future studies with Ussing chambers and patch-clamp techniques.

Contrary to earlier data [13], *ZnT1* gene expression was not regulated in any gut tissue examined. This protein is located on the basolateral side of enterocytes and transports cytosolic Zn into the circulation [11]. *ZnT1* transcription reacts to free cytosolic Zn^2+^, which activates metal-regulatory transcription factor 1 (MTF1) with rising concentrations [13, 20, 21]. Although total jejunal and colonic Zn have been decreased under the present experimental conditions[36], this did not seem to affected the free cytosolic Zn^2+^ according to the unchanged *ZnT1* transcription. Given the aforementioned decrease in jejunal and colonic Zn, it can be speculated that the ZnT1 transport activity was altered, but regulation may have occurred mainly at a post-transcriptional level. The exact nature of this regulation has yet to be investigated. It is assumed that extending the experimental phase >8 d would also have caused an adaption of the *ZnT1* gene expression, as has been shown under long-term conditions [11].

Previous studies showed no significant reaction of kidney or urinary Zn to varying Zn feeding [26, 67]. In fact, the importance of the kidney in maintaining whole-body Zn homeostasis tends to be underestimated. The stabilization of Zn levels in a tissue and its excretions despite changing dietary conditions is, however, a clear sign of active homeostatic regulation. Most previous studies conducted experiments for ≥2 weeks. Hence, the published kidney data reflect an endpoint in adjusting to clinical Zn deficiency. Under the present conditions, kidney Zn and bone Zn showed a strong correlation (r = +0.91, P = 0.002, data not shown). The kidney seemed to give up its Zn in favor of other tissues (e.g. heart, immune tissue) [36]. The present study complements this picture with the reponse patterns of Zn transporter genes including *ZnT1* and *ZIP4*. The first appeared to be regulated in context to basal Zn homeostasis, given its breakpoint at ~60 mg/kg diet. This regulation pattern significantly correlates to kidney Zn and bone Zn (r = +0.77 and 0.94, P = 0.02 and 0.0005, data not shown), which could indicate that ZnT1 was responsible for the kidney Zn depletion in response to the gradual exhaustion of bone Zn. This is in line with earlier reports on the presence of ZnT1 at the basolateral membrane of kidney cells [12]. It further indicates that the renal free cytosolic Zn^2+^ levels correlated with total kidney Zn over a wide range of dietary Zn dosages (28 to 68 mg Zn/kg diet), which contrasts our findings from the intestines. Contrary to the renal *ZnT1* expression, *ZIP4* expression changed at a much lower dietary threshold (~40 mg/kg) and increased linearly with further decrease in dietary Zn. As mentioned above, this breakpoint may indicate the *ZIP4* expression was regulated to compensate for increased cellular stress. An earlier study confirmed ZIP4 gene expression in the kidney [8], but there is yet no data on its role or regulation in renal tissue. We suspect that ZIP4 is involved in recycling of Zn from primary urine under SZD conditions. This would be particularly interesting considering that *ZIP10* gene expression was absent in this tissue, although it was previously claimed to be involved in urinary Zn reabsorption [54–56]. Unfortunately, our data do not allow us to differentiate between kidney cell types. Future studies will need to use microdissections to further support our findings. The possible interaction of the kidney with the intestinal-pancreatic axis of Zn homeostatic regulation, as suggested by Liuzzi, Bobo [7], should also be clarified.

Other Zn transporter genes also showed a significant dose-response to changes in dietary Zn and were shown accordingly in result tables and figures. Their exact role in the examined tissues is currently unclear and requires further scientific studies. In contrast, some *ZnT* and *ZIP* genes showed a very stable expression level over dietary Zn dosages, including jejunal *ZnT5* and *ZnT6*, colonic *ZIP1* and *ZnT5*, hepatic *ZIP7* and *ZIP13* as well as nephric *ZnT4, ZnT5* and *ZnT6*. Therefore, these served as reference genes for data normalization. Future studies should investigate under what conditions the transcription of these genes changes in pigs.

In conclusion, we recognized significant differences in the expression of zinc transporter genes to SZD compared to previous studies on clinical Zn deficiency. Many of the transcripts examined showed significant breakpoints in response to a reduction in dietary Zn. These thresholds were either ~40 or ~60 mg Zn/kg diet, which suggests differences in the specific stimuli to which these genes respond. A breakpoint near ~60 mg Zn/kg diet corresponds to the gross Zn requirement threshold under the present experimental conditions and suggests a role of certain genes in the maintenance of the basal whole-body Zn homeostasis. Other genes showed breakpoints close to ~40 mg Zn/kg diet, which was previously associated with replenishment of tissue Zn and the adjustment of compensation mechanisms in response to increased oxidative stress under SZD conditions. This manuscript presents the first comparative study of the effects of short-term finely-graded differences in dietary Zn supply on the expression of all known ZnT and ZIP genes in different tissues of weaned piglets. Future studies must address their regulation on the proteomic level, accompanied by studies on tissue transcriptomics, proteomics and metabolomics to complete our knowledge on Zn transporters in mammal biology. Our current findings can be translated from pigs into humans due to the high similarity between these species in terms of their nutrition physiology.

## CRediT author statement

**Brugger:** Conceptualization, Methodology, Validation, Formal Analysis, Visualization, Project administration, Writing – Original Draft Preparation, Reviewing and Editing **Hanauer:** Investigation **Ortner:** Investigation **Windisch:** Resources, Writing – Reviewing and Editing, Supervision, Funding Acquisition.

## Acknowledgements

We would like to express our deepest gratitude to Dipl. Ing. (FH) Michael Gertitschke and Andrea Reichlmeir for excellent technical assistance. Special thanks are related to Dr. med. vet. Astrid Kunert for veterinary assistance and advice as well as Gavin Boerboom, M.Sc., and Brett Boden, M.Sc., for valuable advice on the manuscript.

---

**Supplementary Table 1.**
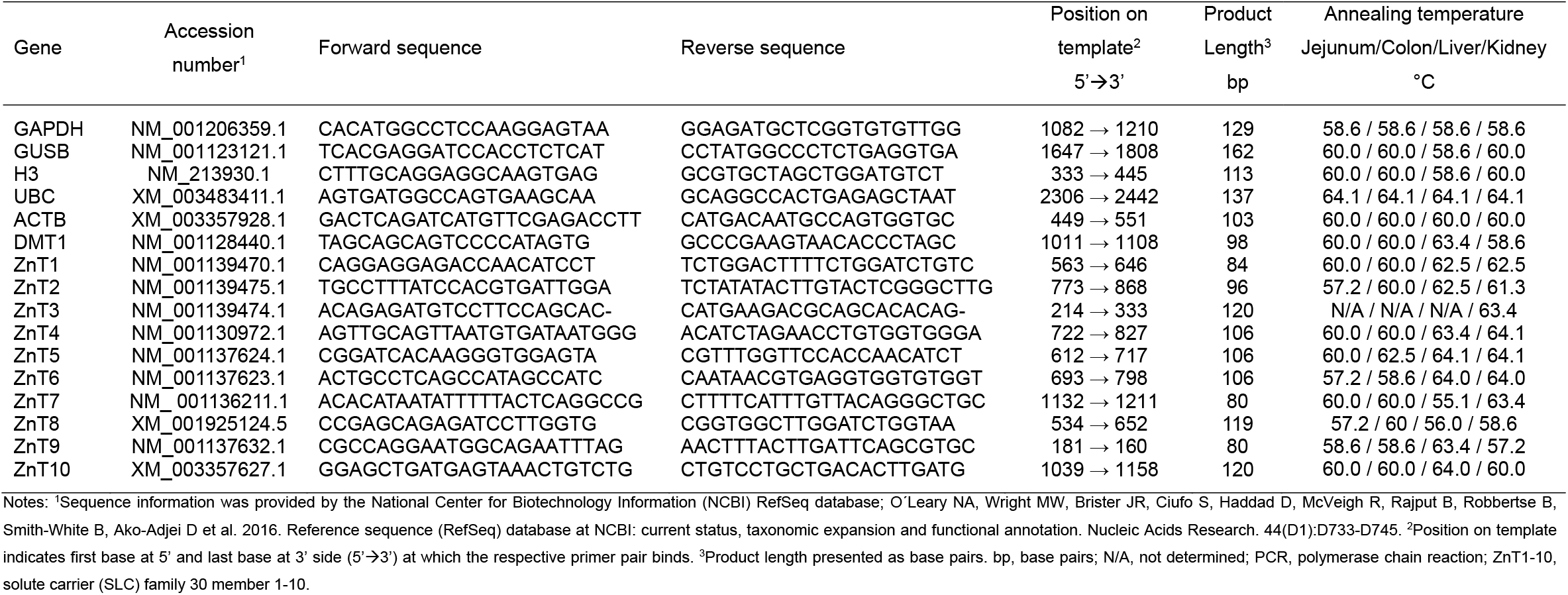
PCR primer and PCR product specifications – Part I

**Supplementary Table 2.**
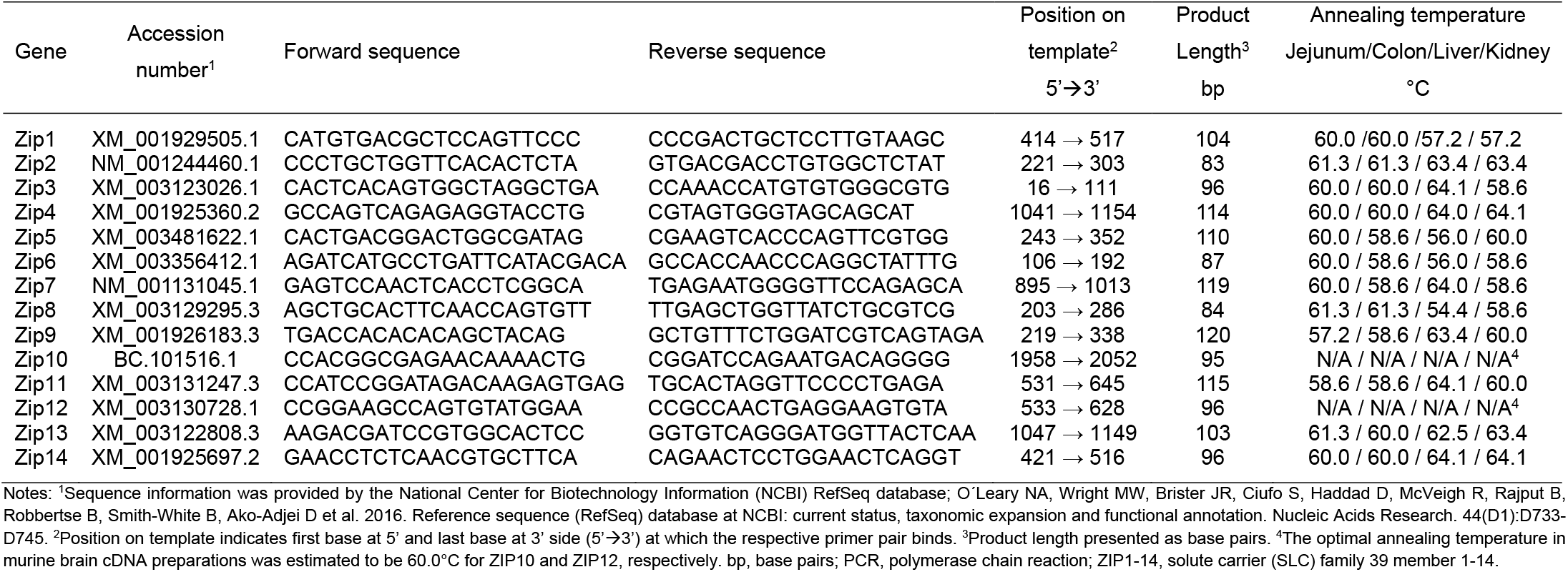
PCR primer and PCR product specifications - Part II

**Supplementary Figure 1.**
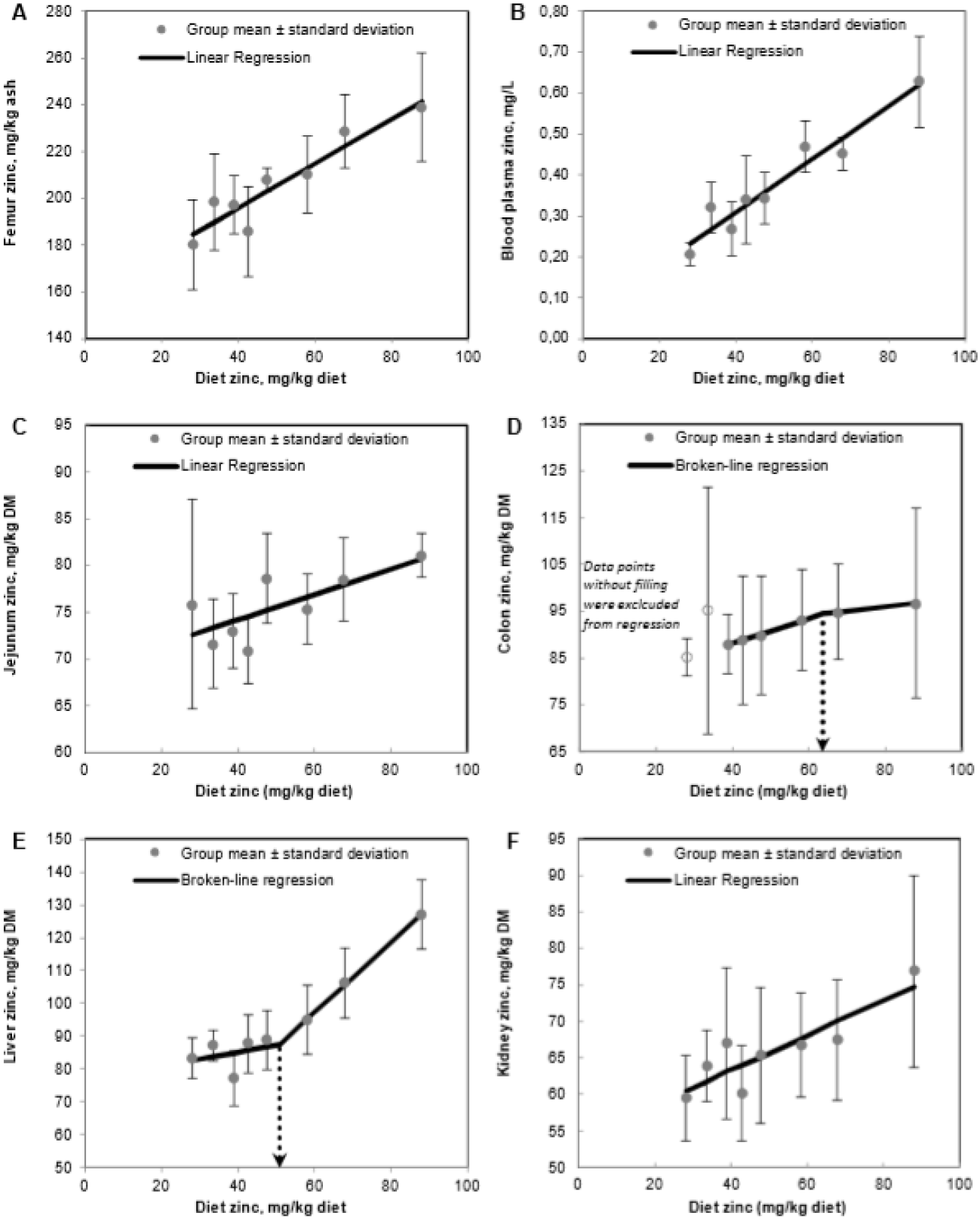
Changes in total zinc pools of femur (A), blood plasma (B), jejunum (C), colon (D), liver (E) and kidney (F) of weaned piglets fed varying dietary Zn for 8 d. Data was originally presented by *Brugger, D., and W. M. Windisch. 2019. Brit. J. Nutr. 121: 849-858*. d, days; DM, dry matter; Values are arithmetic means ± SDs, n = 6 single values per mean.

